# A clinical-anatomical signature of Parkinson’s Disease identified with partial least squares and magnetic resonance imaging

**DOI:** 10.1101/168989

**Authors:** Yashar Zeighami, Seyed-Mohammad Fereshtehnejad, Mahsa Dadar, D. Louis Collins, Ronald B. Postuma, Bratislav Mišić, Alain Dagher

## Abstract

Parkinson’s disease (PD) is a neurodegenerative disorder characterized by a wide array of motor and non-motor symptoms. It remains unclear whether neurodegeneration in discrete loci gives rise to discrete symptoms, or whether network-wide atrophy gives rise to the unique behavioural and clinical profile associated with PD. Here we apply a data-driven strategy to isolate large-scale, multivariate associations between distributed atrophy patterns and clinical phenotypes in PD. In a sample of N = 229 de novo PD patients, we estimate disease-related atrophy using deformation based morphometry (DBM) of T1 weighted MR images. Using partial least squares (PLS), we identify a network of subcortical and cortical regions whose collective atrophy is associated with a clinical phenotype encompassing motor and non-motor features. Despite the relatively early stage of the disease in the sample, the atrophy pattern encompassed lower brainstem, substantia nigra, basal ganglia and cortical areas, consistent with the Braak hypothesis. In addition, individual variation in this putative atrophy network predicted longitudinal clinical progression in both motor and non-motor symptoms. Altogether, these results demonstrate a pleiotropic mapping between neurodegeneration and the clinical manifestations of PD, and that this mapping can be detected even in *de novo* patients.

## 1. Introduction

Parkinson’s disease (PD) is a neurodegenerative disease characterized by progressive and widespread neuronal loss associated with intracellular aggregates of α-synuclein giving rise to the classical Lewy pathology (Goedert et al., 2013; Poewe et al., 2017). PD has been traditionally known as a motor disease with bradykinesia, rigidity, and tremor as the cardinal symptoms, and preferential loss of dopamine neurons of the substantia nigra. The motor symptoms have been the main target for diagnosis and treatment (Kalia and Lang, 2015). However, it is now clear that PD is a more complex disorder involving several non-motor manifestations that both precede and follow the initial appearance of motor symptoms. The non-motor aspects of PD involve several clinical domains including autonomic, limbic, olfactory, and cognitive (Chaudhuri et al., 2006; Poewe, 2008). A 15-year follow-up study shows cognitive decline and dementia in up to 80% of surviving PD patients (Hely et al., 2005).

Over time, PD diagnostic criteria have been modified toward a multifaceted characterization in response to the insufficiency of the narrow motor definition of PD (Postuma et al., 2016). The increasing attention to non-motor aspects of the disease has allowed detection of more diverse clinical patterns in PD. For example, recent studies have subcategorized PD patients based on the dominance of motor, rapid eye movement sleep behavior disorder (RBD), autonomic, and cognitive deterioration (Fereshtehnejad et al., 2015; Fereshtehnejad et al., 2017). Post-mortem and neuroimaging studies have emphasized the preferential loss of dopamine neurons in the substantia nigra (Halliday and McCann, 2010). However, post-mortem studies have also shown that the pathological process is neither initiated in nor confined to the substantia nigra, gradually ascending from the olfactory tracts and medulla to the midbrain and cortical layers (Braak et al., 2003; Goedert et al., 2013).

Neuroimaging studies in PD have evolved in the past 30 years (Politis, 2014). The main focus of early studies was on dopaminergic innervation, using single photon emission computed tomography (SPECT) or positron emission tomography (PET). However, the availability of new higher resolution whole-brain neuroimaging techniques such as magnetic resonance imaging (MRI), metabolic imaging with ^18^F-FDG PET, and resting or task state functional MRI have provided the opportunity to investigate the non-dopaminergic aspects of PD (Politis, 2014; Tuite and Dagher, 2013; Yousaf et al., 2017). Structural analysis using MRI (including T1, T2, and diffusion weighted MRI) was initially inconclusive, or only sensitive enough to capture disease related differences in late stages of PD once dementia had set in. More recently, with larger sample sizes and higher resolution imaging, it has been possible to study *de novo* PD patients using MRI (Heim et al., 2017; Zeighami et al., 2015). However, these studies mostly focus on brain related differences between PD and healthy control populations, or on a single aspect of the disease (e.g. dementia or motor symptoms) as post hoc analysis. To our knowledge, no studies have attempted to model the relationship between brain atrophy and presence and severity of the entire constellation of motor and non-motor symptoms in PD simultaneously. Such an approach might also make it possible to disambiguate different domains or modes of the disease within one PD population and their relationship with brain morphometric measures.

Here we use a multivariate method to relate the motor and non-motor aspects of PD to systemwide atrophy patterns. We use data from 235 newly diagnosed PD patients and 117 age- and sex-matched healthy controls from the Parkinson’s Progression Markers Initiative (PPMI) database (www.ppmi-info.org/data), an observational, multicenter longitudinal study designed to identify PD progression biomarkers (Marek et al., 2011). We use deformation-based morphometry (DBM), which is based on local nonlinear subject-to-template deformations as a measure of structural brain alterations (Ashburner et al., 2000; Aubert-Broche et al., 2013; Chung et al., 2001; Penny et al., 2011), and partial least squares (PLS) (McIntosh and Lobaugh, 2004; McIntosh and Misic, 2013; Wold, 1966) to capture the relationship between brain atrophy patterns and disease-related clinical measures. Furthermore, we explore the extent to which brain atrophy patterns can predict disease progression by examining longitudinal changes across different measures of disease severity.

## 2. Methods

### 2.1. PPMI dataset

Data used in the preparation of this article were obtained from the Parkinson’s Progression Markers Initiative (PPMI) database (http://www.ppmi-info.org/data). For up-to-date information on the study, see www.ppmi-info.org. PPMI is a cohort of people with *de-novo* idiopathic PD (Marek et al., 2011). Individuals were eligible for recruitment if they were at least 30 years old, diagnosed with PD within the last 2 years, had at least two signs or symptoms of Parkinsonism (tremor, bradykinesia and rigidity), a baseline Hoehn and Yahr Stage of I or II, and did not require symptomatic treatment within six months of the baseline visit. The PPMI is a multi-center international project and the institutional review boards approved the protocol at all participating sites. Participation was voluntary and all individuals signed the written informed consent prior to inclusion.

We obtained data from the baseline visit 3T high-resolution T1-weighted MRI scans in compliance with the PPMI Data Use Agreement. For clinical data, any participant with > 20% missing values at baseline was excluded. Overall, MRI and clinical data were included for 229 drug-naïve participants with PD (6 subjects failed MRI quality control) and 117 healthy sex- and age-matched controls (78 male, age = 59.2 ± 11.3 years). All subjects including patients and healthy controls were part of the PPMI dataset. Healthy controls were eligible if they were aged >30 years, had no history of neurological disease and no first degree relative with PD (Marek et al., 2011).

For each subject, we also obtained demographic and clinical information as well as cerebrospinal fluid (CSF) and SPECT biomarker values from the dataset in May 2016 (accession date). General information consisted of age at disease onset, gender, years of education, handedness and disease duration. Clinical and laboratory markers are described below.

### 2.2. Brain Imaging Data Analysis

MRI data consisted of 1×1×1 mm 3T T1-weighted scans obtained from the PPMI database. All scans were pre-processed through an in-house MR image processing pipeline, using image de-noising (Coupe et al. 2008), intensity non-uniformity correction (Sled et al., 1998), and image intensity normalization using histogram matching. The preprocessed images were first linearly (using a 9-parameter rigid registration) and then nonlinearly registered to a standard brain template (MNI ICBM152) (Collins and Evans, 1997; Collins et al., 1994). Using the obtained nonlinear transformations, deformation based morphometry (DBM) was performed to calculate local density changes as a measure of tissue expansion or atrophy. For more detail on the processing steps please see Zeighami et al. (2015). We obtained a single deformation brain map for each subject. The value at each voxel is equal to the determinant of the Jacobian of the transformation matrix obtained from nonlinear registration of participants’ T1 MR images and MNI-ICBM152 brain template. The DBM values reflect regional brain deformations and can be used as indirect measures of brain atrophy (Cardenas et al., 2007; Chung et al., 2001; Leow et al., 2006; Studholme et al., 2004).

### 2.3. Clinical Measures

PD-related motor, cognitive and non-motor clinical manifestations were assessed at baseline and each follow-up visit (Table 1).

**Table 1.**
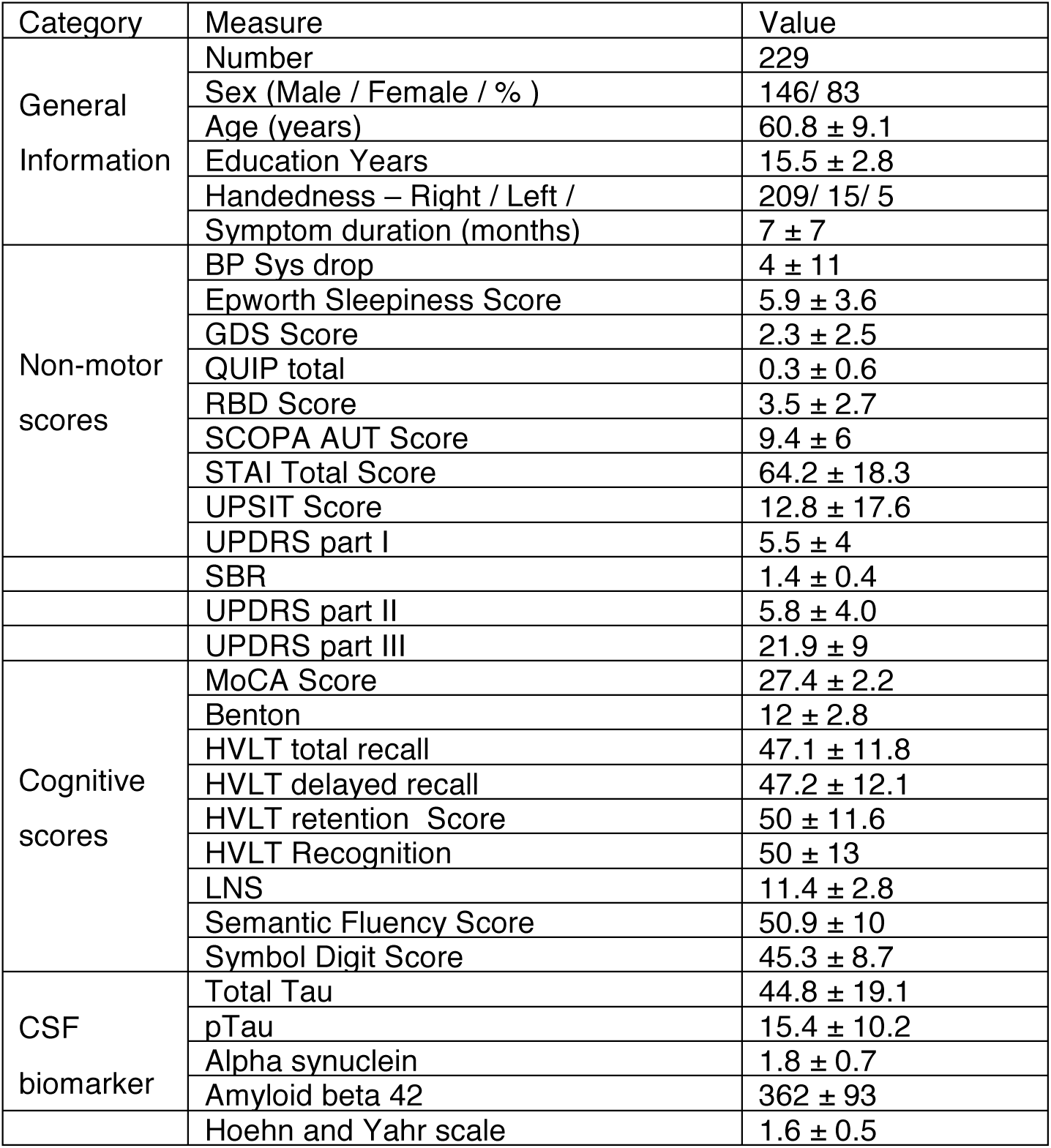
Demographic and clinical information for individuals with Parkinson’s disease from the PPMI used in this study. BP Sys= Systolic Blood Pressure. GDS= Geriatric Depression Scale. QUIP = Questionnaire for Impulsive-Compulsive Disorders. RBD= REM sleep behaviour disorder. SCOPA= Scales for Outcomes in PD-Autonomic. STAI= State-Trait Anxiety Inventory. UPDRS= Unified Parkinson’s Disease Rating Scale. SBR= striatal binding ratio. MoCA= Montreal Cognitive Assessment. HVLT= Hopkins Verbal Learning Test. LNS= letter-number sequencing. All error terms used are standard deviations.

We also included Genetic Risk Score in the PLS analysis. This is a single surrogate indicator that summarizes 30 risk alleles for PD (Nalls et al., 2015). All clinical assessments were repeated in follow-up visits (minimum= 1 year, mean= 2.7 years). In order to evaluate disease progression, we created a putative global composite outcome (GCO) as a single indicator by combining z-scores of the most clinically relevant motor and non-motor measures of disease severity including Movement Disorder Society Unified Parkinson’s Disease Rating Scale (MDS-UPDRS) parts I, II, and III, Schwab and England activities of daily living (SE-ADL) score, and Montreal Cognitive Assessment (MoCA) score as described previously (Fereshtehnejad et al., 2017).

### 2.4. Biomarkers

The striatal binding ratio (SBR), a marker of dopaminergic denervation in caudate and putamen, was obtained by SPECT with the DAT tracer ^123^I-Ioflupane at baseline and follow-up. Cerebrospinal fluid (CSF) biomarkers consisting of amyloid-beta (Aβ1-42), total Tau (T-tau), phosphorylated tau (P-tau181) and α-synuclein were also included in our analysis. Information for all variables is summarized in Table 1.

### 2.5. Partial least squares analysis

Partial least squares (PLS) is an associative, multivariate method for relating two sets of variables to each other (Abdi and Williams, 2010; McIntosh and Lobaugh, 2004; McIntosh and Misic, 2013; Wold, 1966). The analysis seeks to find weighted linear combinations of the original variables that maximally covary with each other. Here, the two variable sets were voxel-wise brain atrophy (as measured by DBM) and clinical/demographic measures (Table 1). The respective linear combinations of these variables can be interpreted as atrophy networks and their associated clinical phenotypes.

#### Singular value decomposition

The imaging and clinical data were organized in two matrices, **X** (DBM) and **Y** (clinical), with participants in the rows of the matrices and variables in the columns (Figure 1). Both matrices were first z-scored by subtracting the mean from each column (variable) and dividing by the standard deviation. The atrophy-clinical covariance matrix was then computed, representing the covariation of all voxel deformation values and clinical measures across participants. Since the data are z-scored, the atrophy-clinical covariance is effectively a correlation matrix. The resulting matrix was then subjected to singular value decomposition (SVD) (Eckart and Young, 1936):

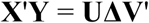

such that

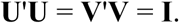

**Figure 1.**
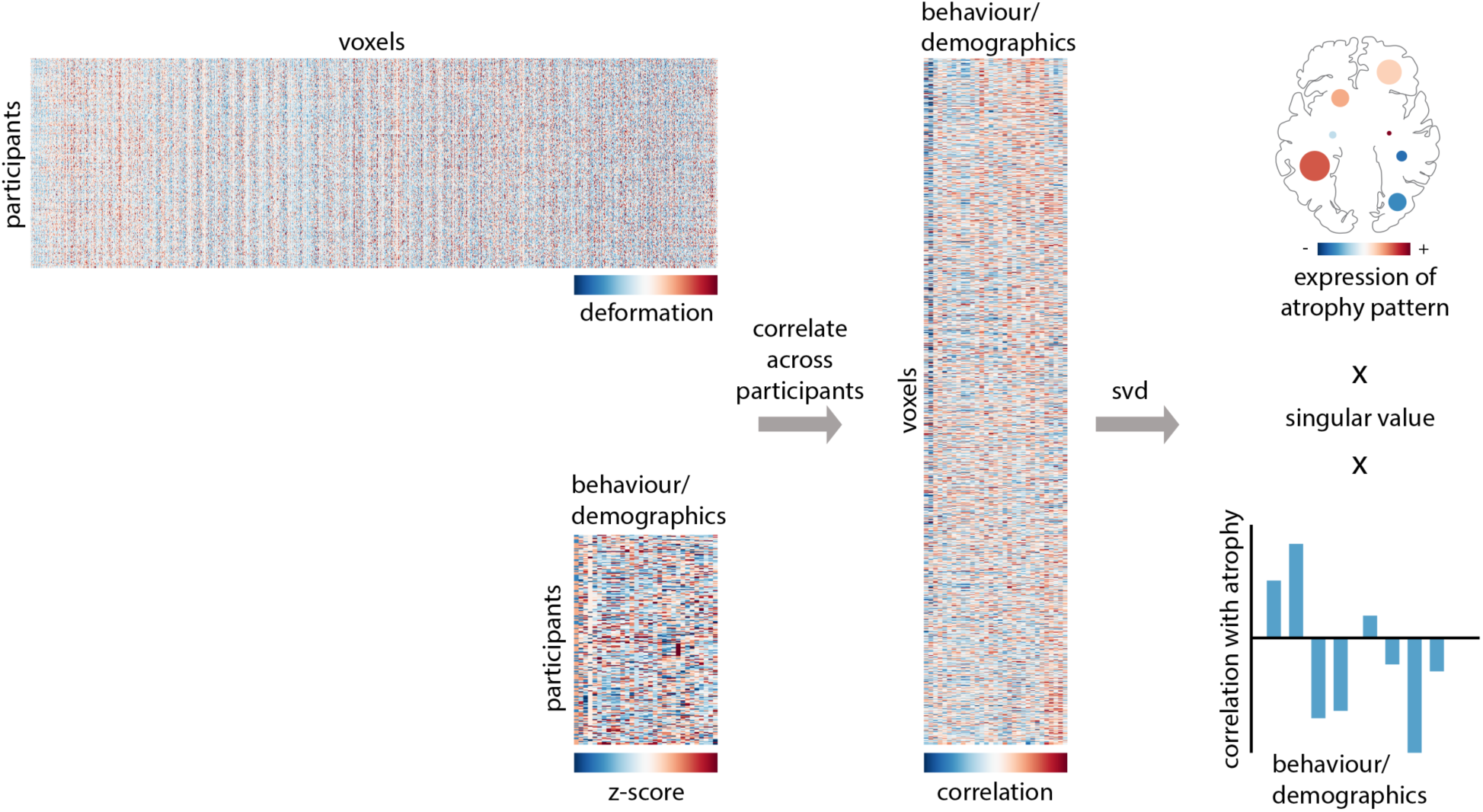
Partial Least Square (PLS) Analysis flowchart.

The decomposition yields a set of mutually orthogonal latent variables (LVs), where **U** and **V** are matrices of left and right singular vectors, and **Δ** is a diagonal matrix of singular values. Each latent variable is a triplet of the i^th^ left singular vector, the i^th^ right singular vector and the i^th^ singular value. The number of latent variables is equal to the rank of the covariance matrix, which is the smaller of its dimensions or the dimension of its constituent matrices. In the present study, the number of clinical measures (k = 31) is the smallest dimension, so the rank of the matrix and the total number of latent variables is equal to 31. If there are v voxels, the dimensions of **U**, **V**, and Δ are v × k, k × k, and k × k, respectively.

Each singular vector weights the original variables in the multivariate pattern. Thus, the columns of **U** and **V** weight the original voxel deformation values and clinical measures such that they maximally covary. The weighted patterns can be interpreted as a set of maximally covarying atrophy patterns and their corresponding clinical phenotypes. Each such pairing is associated with a singular value from the diagonal matrix, proportional to the covariance between atrophy and behavior captured by the latent variable. Specifically, the effect size associated with each latent variable (proportion of covariance accounted for) can be naturally estimated as the ratio of the squared singular value to the sum of all squared singular values (McIntosh and Lobaugh, 2004).

#### Significance of multivariate patterns

The statistical significance of each latent variable was assessed by permutation tests. The ordering of observations (i.e. rows) of data matrix **X** was randomly permuted (N = 500 repetitions), and a set of “null” atrophy-behavior correlation matrices were then computed for the permuted brain and non-permuted clinical data matrices. By permuting the order of patients, the procedure effectively destroys any dependencies between atrophy and behavior. These “null” correlation matrices were then subjected to SVD as described above, generating a distribution of singular values under the null hypothesis that there is no relationship between brain deformation and clinical measures. Since singular values are proportional to the magnitude of a latent variable, a non-parametric P value can be estimated for a given latent variable as the probability that a permuted singular value exceeds the original, nonpermuted singular value. Of note, the permutation test generates a composite set of p-values from a single multivariate test, implicitly embodying control of type II error.

#### Contribution and reliability of individual variables

The contribution of individual variables (voxels or clinical measures) was estimated by bootstrap resampling. Participants (rows of data matrices **X** and **Y**) were randomly sampled with replacement (N = 500), generating a set of resampled correlation matrices that were then subjected to SVD. This procedure generated a sampling distribution for each individual weight in the singular vectors. A “bootstrap ratio” was calculated for each voxel as the ratio of its singular vector weight and its bootstrap-estimated standard error. Thus, large bootstrap ratios can be used to isolate voxels that make a large contribution to the atrophy pattern (have a large singular vector weight) and are stable across participants (have a small standard error). If the bootstrap distribution is approximately normal, the bootstrap ratio is equivalent to a z-score (Efron and Tibshirani, 1986). Bootstrap ratio maps were thresholded at values corresponding to the 95% confidence interval.

#### Patient-specific atrophy and clinical scores

To estimate the extent to which individual patients express the atrophy or behavioural patterns derived from the analysis, we calculated patient-specific scores. Namely, we projected the weighted patterns **U** and **V** onto individual-patient data, yielding a scalar atrophy score and clinical score for each patient, analogous to a principal component score or factor score:

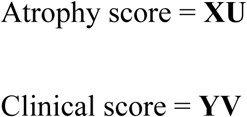

To investigate the predictive utility of the PLS model, we correlated patient-specific atrophy and clinical scores with longitudinal measures of disease progression. These included the Global Composite Outcome (GCO) and SE-ADL scores as measures of general disease severity, MoCA for cognition, MDS-UPDRS III for motor, and MDS-UPDRS I for non-motor aspects of disease.

## 3. Results

### 3.1. PLS analysis

The PLS analysis revealed six statistically significant latent variables relating clinical measures in PD and their corresponding brain atrophy patterns (permuted p <0.0001, p<0.005, p<0.05, p<0.05, p< 0.005, p<0.05). These patterns respectively account for 17.5, 9, 8.2, 6, 4.6, and 4.5% (total of 50%) of the shared covariance between clinical and brain atrophy measures. Based on the variance explained and clinical interpretability of the results, we focus on and discuss the first latent variable (LV-I) in greater detail (Figure 2).

**Figure 2.**
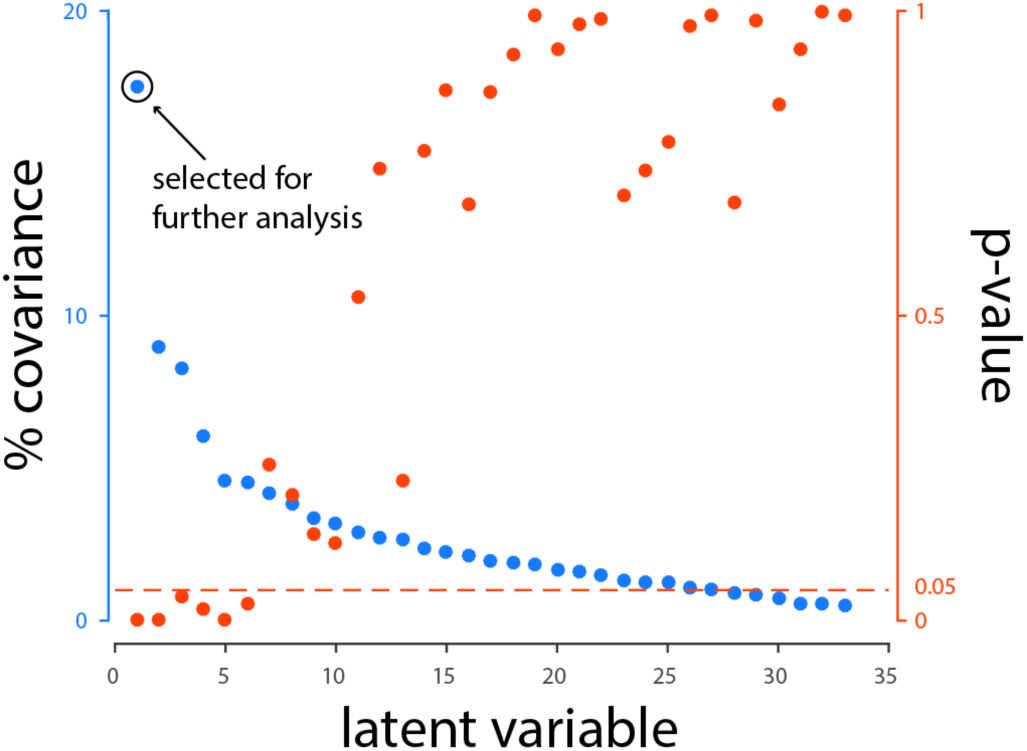
Covariance explained and permutation p-values for all latent variables in the PLS analysis. LV-I is selected for further analysis based on the variance explained and clinical interpretability of the results. PLS = Partial Least Squares.

### 3.2. Clinical features and biomarkers patterns

The biomarkers and clinical features (Figure 3) contributing to LV-I are composed of: higher PD-related severity (motor and non-motor) as measured by UPDRS scores, lower striatal dopamine innervation measured by SPECT, lower cognitive performance (mainly memory-related), lower amyloid beta level in CSF, and more severe anxiety, depression, and sleep disorder. We also found the previously reported effects of age (worse with age) and gender (males worse). More specifically, age was the strongest contributor to LV-I (R = 0.69, 95% CI [0.59,0.74]) followed by motor signs measured by UPDRS-III (R = 0.35, 95% CI [0.35,0.52]) and autonomic disturbances (SCOPA-AUT) (R = 0.27, 95% CI [0.25,0.45]). Male gender (R = 0.23, 95% CI [0.07,0.34]) and symptom duration (R = 0.19, 95% CI [0.10,0.32]) were other significant contributors to LV-I. Impaired visuospatial (Benton Line Orientation) (R = -0.25, 95% CI [-0.40,-0.18]) and executive function (Letter-Number Sequencing) (R = -0.25, 95% CI [-0.38,-0.16]) were the strongest cognitive features of LV-I, followed by the global cognitive status measured by MoCA (R = -0.21, 95% CI [-0.35,-0.12]), impaired speed/attention domain (Symbol-Digit Matching) (R = -0.19, 95% CI [-0.35,-0.11]) and memory deficit (HVLT) (R = -0.13, 95% CI [-0.28,-0.05]). CSF concentration of amyloid-beta (R = -0.17, 95% CI [-0.31,-0.03]) and severity of dopaminergic denervation (SBR) (R = -0.15, 95% CI [-0.32,-0.06]) were the only biomarkers that significantly contributed to LV-I, while genetic risk score (R = -0.09, 95% CI [-0.23,0.04]) and CSF α-synuclein (R = -0.01, 95% CI [-0.15,0.14]) failed to reach significance levels.

**Figure 3.**
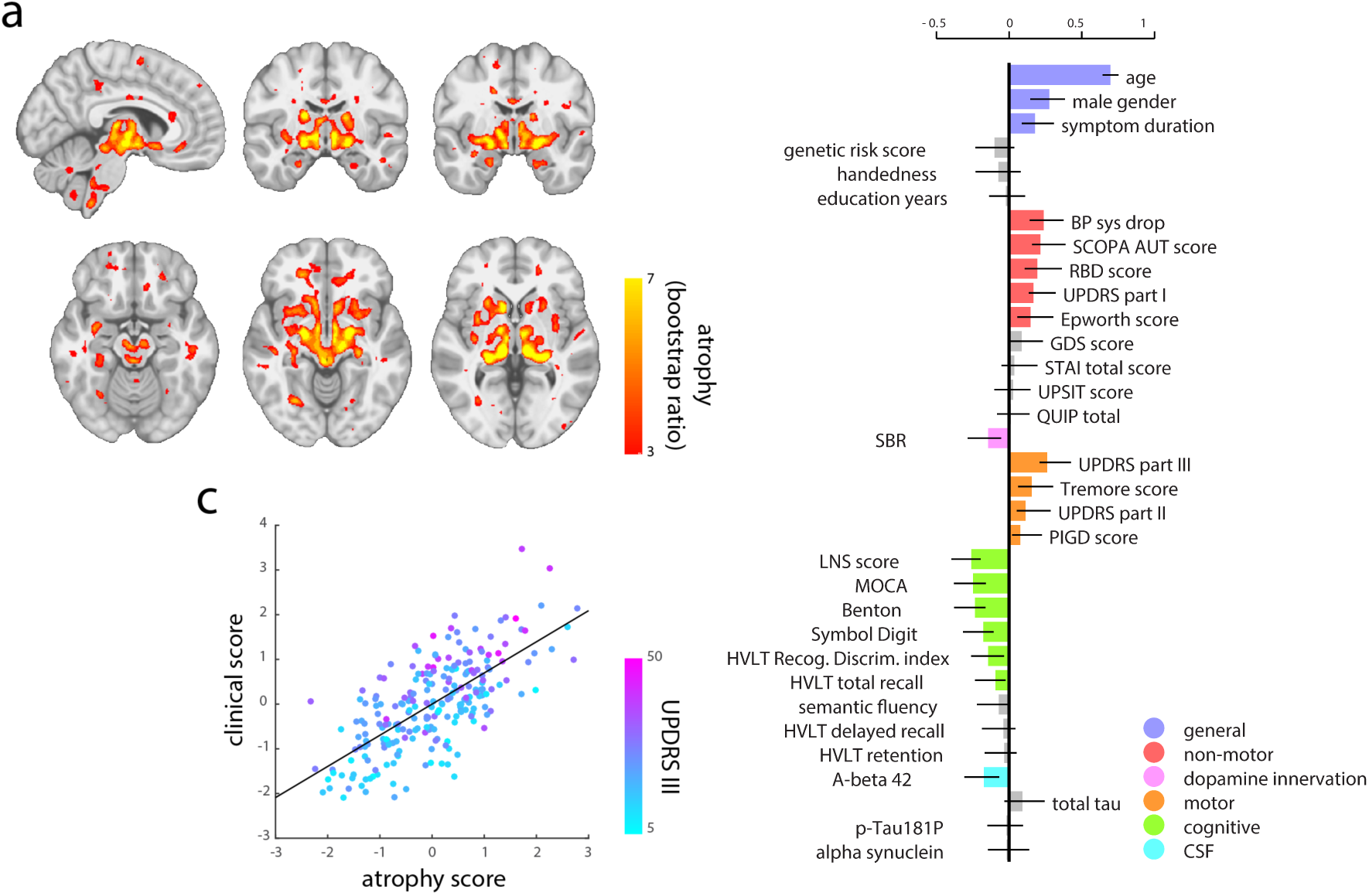
First latent variable (LV-I) obtained from the PLS analysis. a) Brain pattern bootstrap ratios in MNI space (x= -6, y = -12, y= -8, z= -14, z= -6, z= 0) b) Clinical scores pattern. The effect size estimates are derived from SVD analysis and the Confidence Intervals (CI) are calculated by bootstrapping, hence the CI are not necessarily symmetrical. c) individual subjects’ Brain versus Clinical PLS score.

LV-II to LV-IV are shown in the supplementary materials (supplementary figures 1-3). Briefly, LV-II (supplementary figure 1) represents features of a more benign phenotype of PD with a higher tremor score, higher dopamine innervation as measured by SPECT, better memory function (measured by HVLT total recall), and less severe behavioral symptoms, sleep disorders, and autonomic disturbances. By contrast, LV-III (supplementary figure 2) indexes more prominent postural and gait disabilities (PIGD Score) and more severe mood and behavioral symptoms, RBD, and autonomic disturbances. This LV also includes impaired visuospatial cognitive functions and more severe hyposmia as measured by the UPSIT.

### 3.3. Atrophy network in de novo PD patients

The corresponding brain pattern for the clinical and demographic measures in LV-I involved discrete cortical regions located in multiple parts of the frontal lobes, fusiform gyrus, cingulate gyrus and insular cortex, and subcortical regions including thalamus and basal ganglia (putamen, caudate, and nucleus accumbens), hippocampus and amygdala, brainstem (substantia nigra, red nucleus, subthalamic nucleus, pons, and areas of medulla that overlap with the dorsal motor nucleus of the vagus and nucleus of the solitary tract), and cerebellum. (Table 2, Figure 3.a.) Figure 3c shows an example of how the putative atrophy network and the associated clinical phenotype relate to each other. For each weighted pattern, we estimated patient-specific scores by projecting the patterns onto individual patients’ data (see *Methods*). The resulting scalar values (termed atrophy scores and clinical scores), reflect the extent to which an individual patient expresses each pattern. By definition, the two scores are correlated (r = 0.7), i.e. patients with greater atrophy in the network in Fig. 3a, also tend to conform more closely to the clinical phenotype in Fig. 3b. Patients who score highly on both likely have more severe pathology, and we illustrate this by coloring the points (individual patients) by their UPDRS III scores. Individuals with more pronounced atrophy and clinical variable severity also tend to score highly on UPDRS III, a measure of motor symptoms.

**Table 2.** Peak coordinates in MNI-ICBM152 space for brain PLS scores using bootstrap ratios. PLS = Partial Least Squares. B-ratio= Bootstrap ratio. MNI: Montreal Neurological Institute. Structures are ordered within regions by z coordinate.

| Region | B-ratios | Structure | MNI coordinates |
| --- | --- | --- | --- |
| Brainstem | 4.9 | Medulla | -6,-42,-58 |
|  | 3.6 | Pons | -3,-30,-47 |
|  | 4.7/4.5 | Substantia Nigra | 9,-14,-13/-7,-14,-13 |
|  | 5.1/4.4 | Subthalamic nucleus | 8,16,-10/-8,-16,-10 |
| Cerebellum | 5.6/4.8 | Cerebellum | 32,-64,-34 |
|  | 4.8 | Cerebellum | -26,-64,-32 |
| Subcortical | 4.2/4 | Hippocampus | 23,-9,-25/-22,-9,-26 |
|  | 4.1/4.2 | Amygdala | 20,-4,-23/-25,-4,-24 |
|  | 5.6/5.1 | Nucleus Accumbens | 10,12,-6/-8,12,-10 |
|  | 5.7/6.0 | Globus Pallidus Internal Segment | 22,-6,-4/-22,-8,-4 |
|  | 5.4/4.7 | Putamen | 24,12,-6/-24,12,-4 |
|  | 8.6/7.7 | Ventrolateral/Ventroposterior Thalamus | 12,-26,-2/-10,-24,-2 |
|  | 7.3/4.7 | Caudate | 10,14,-2/-10,14,3 |
| Cortical | 3.2 | Fusiform gyrus | 22,8,-48 |
|  | 3.7 | Medial temporopolar region | -22,10,-44 |
|  | 3.4 | Medial/Inferior frontal gyrus | 50,8,-38 |
|  | 3.2/3.3 | Anterior/medial Orbital Gyrus | 23,49,-14/-23,43,-18 |
|  | 6.4/5.8 | Periaqueductal Gray | 6,-32,-12/-4,-32,-12 |
|  | 5.3/2.8 | Fusiform gyrus | 26,-66,-8/-24,-64,-9 |
|  | 3.7 | Fusiform gyrus | -22,-66,-4 |
|  | 5.0 | Inferior Frontal gyrus | -30,32,6 |
|  | 3.4 | Medial frontal gyrus | 44,52,12 |
|  | 3.6 | Lateral occipital cortex | -24,-78,20 |
|  | 3.7 | Parietal Operculum | 70,-30,22 |
|  | 2.6 | Cingulate gyrus | 11,22,31 |
|  | 2.8 | Middle Frontal gyrus | 24,33,32 |
|  | 3 | Superior Frontal gyrus | 26,-4,68 |
|  | 2.7 | Cingulate gyrus | -8,-26,76 |

### 3.4. Atrophy pattern in Parkinson’s disease and aging

Aging is the largest risk factor for both development and progression of PD (Hindle, 2010). PD-related and age-related brain alterations could happen independently; however, it is more likely that the two phenomena interact in PD (Collier et al., 2011). From a modeling and analysis perspective the interdependency between the two factors makes it difficult if not impossible to disentangle the two processes without losing the disease related effect from the data.

Nonetheless, to ensure the disease specificity of the findings, the PLS analysis was repeated after removing the effect of aging from the atrophy maps, by regressing out age effects on deformation calculated based on the healthy subjects in the same dataset (N=117). This analysis was similar to previous studies with confounding age effects in diseased populations (Scahill et al. 2003; Franke et al. 2010; Dukart et al. 2011; Moradi et al. 2015). The brain-clinical relationship as identified using PLS remained significant after controlling for normative aging. Overall, the directionality and significant contributors of the age removed LV (AR-LV) patterns remained similar after regressing out the effect of age. We found eight statistically significant LVs relating clinical measures in PD and their corresponding brain atrophy patterns (AR-LV-I to AR-LV-VI permuted p <0.0001, AR-LV-VII and AR-LV-VIII permuted p<0.05). These patterns respectively account for 12, 11, 8, 6.6, 6, 5, 4.7, 4, and 3.6% (total of 55%) of the shared covariance between clinical and brain atrophy measures (Figure 4).

**Figure 4.**
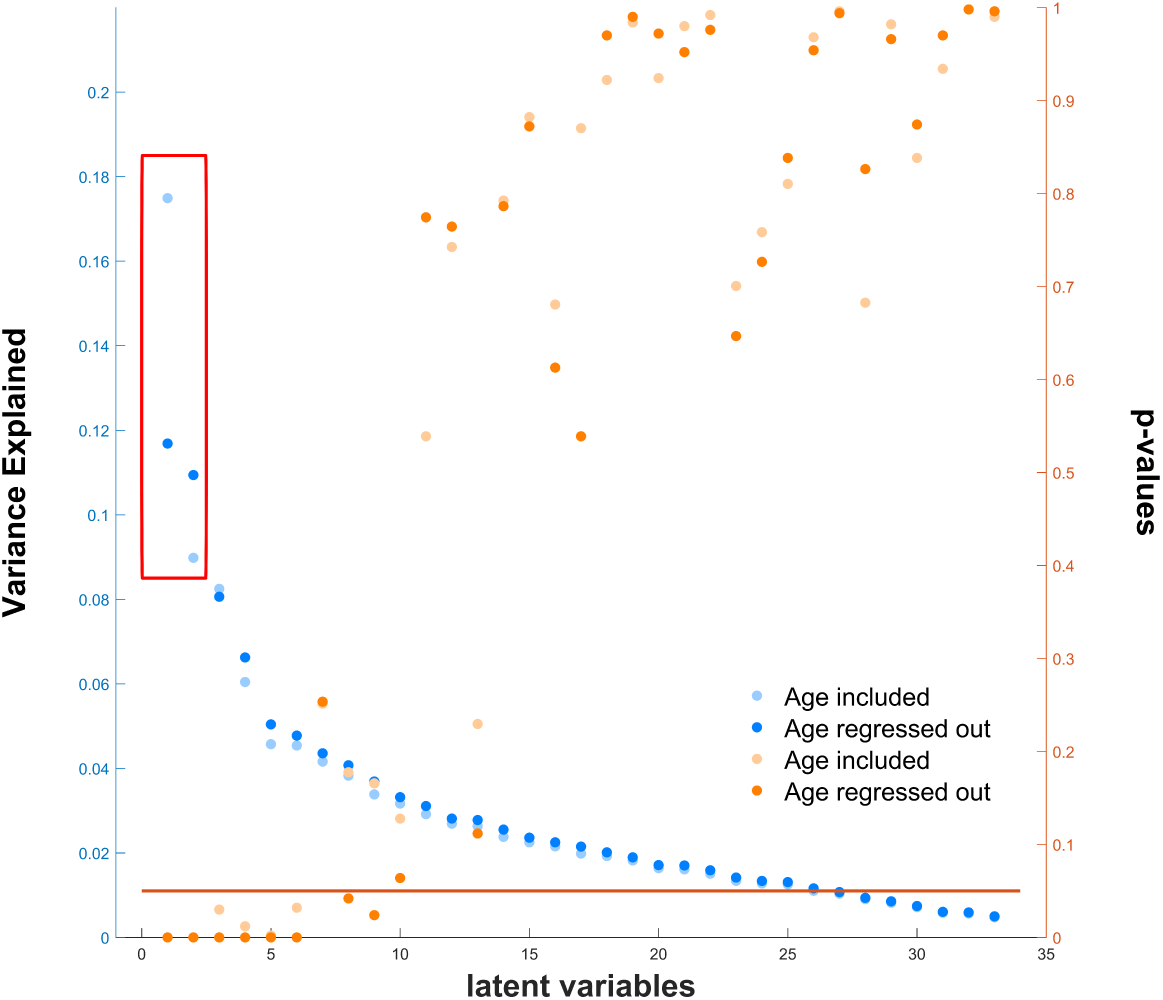
Covariance explained and permutation p-values for all latent variables in PLS analysis before and after regressing out healthy aging in the deformation maps.

Here we focus on the first two LVs since the others remain intact to the effect of regressing out normative aging (Figure 4).

AR-LV-I and AR-LV-II explain more than 20% of the covariance between brain atrophy and clinical measure included in this analysis. AR-LV-I (Figure 5) is similar to LV-I except that age no longer features. It captures the male gender effect, memory-specific cognitive impairment, RBD, as well as certain mood/affective behavioral scores such as the GDS (measuring depression), QUIP (measuring impulse control disorder), and STAI (measuring anxiety disorder) that are absent in LV-I (without controlling for brain normative aging).

**Figure 5.**
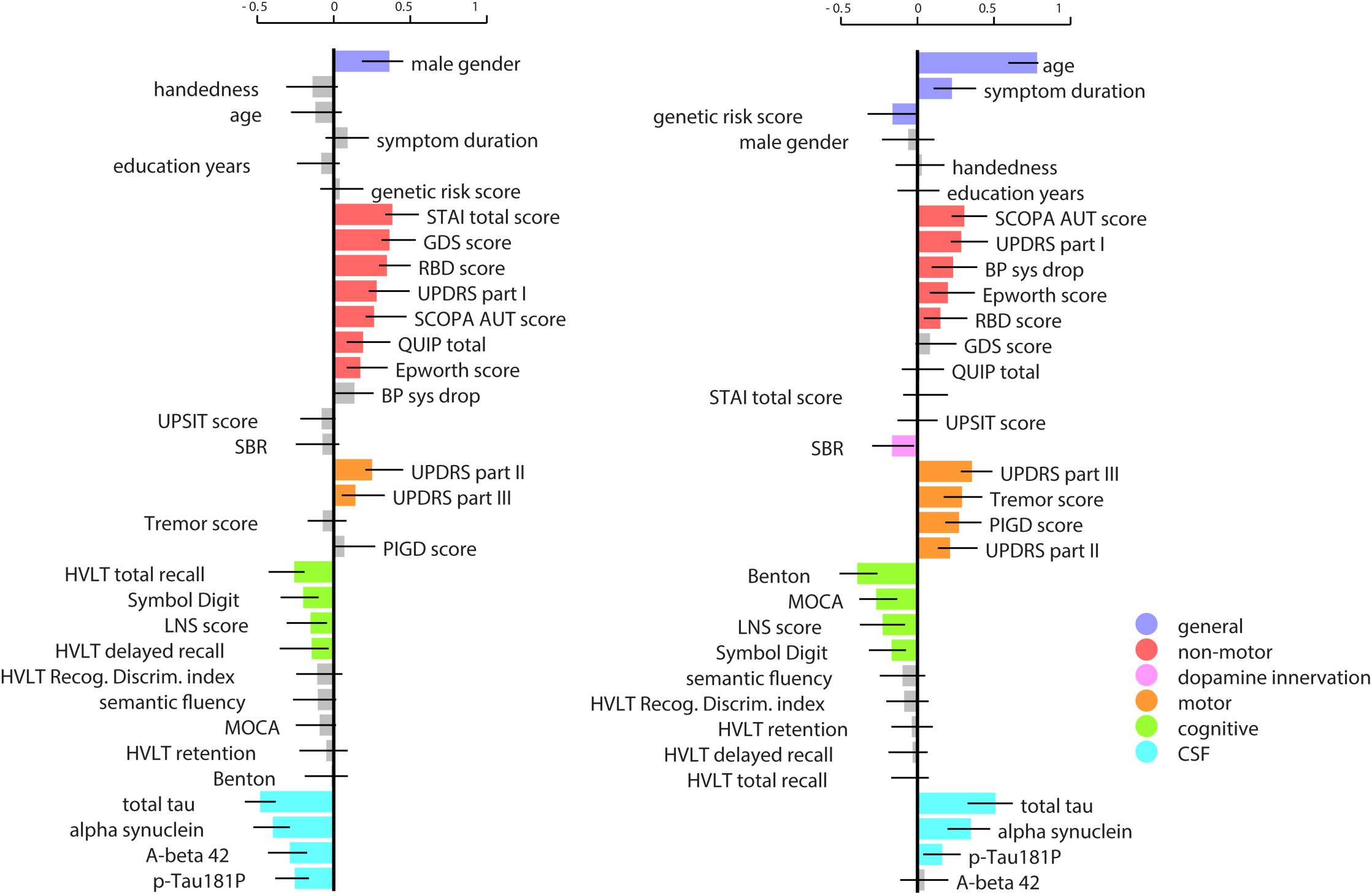
First (left) and second (right) latent variable (AR-LV-I and II) obtained from PLS analysis after regressing out healthy aging. Clinical scores pattern (the effect sizes are estimated using SVD analysis and the Confidence Intervals (CI) are calculated by bootstrapping). PLS=Partial Least Squares. SVD=Singular Value Decomposition.

For AR-LV-II (Figure 5), the significant contributors and their overall directionality are analogous to LV-I, except for CSF measures and gender. The most important contributor in AR-LV-II is age, which suggests that aging contributes to brain alteration in PD beyond the normative aging process. The increase in contribution of motor symptoms (as measured by UPDRS-III) and phenotype (as measured by PIGD) is in line with the significant impact of symptom duration and pathological aging within this LV. In sum, the first two LVs of the age-regressed analysis appear to capture separate portions of the first LV from the non-age-regressed analysis. The AR-LV-III and AR-LV-IV are shown in supplementary figures 4-5.

### 3.5. Atrophy pattern at first visit correlates with longitudinal disease progression

Baseline LV-I score was significantly related to longitudinal worsening in several clinical measures after an average of 2.7 years (Figure 6). Participants with greater expression (atrophy) of the LV-I brain pattern at baseline had significantly greater deterioration in the GCO (r = 0.22, p < 0.001) and in activities of daily living, measured by the SE-ADL (which was not included in the PLS analysis) (r = - 0.20, p=0.003). We also assessed the correlation between LV-I score at baseline and changes in single clinical measures in different categories. Higher expression of the Brain LV-I pattern was significantly correlated with decline in cognition demonstrated by the decrease in MoCA score (r = -0.28, p<0.0001). However, the association between baseline LV-I expression and changes in motor signs (UPDRS-III) (r = 0.13, p = 0.052) or non-motor symptoms (UPDRS-I) (r = 0.12, p=0.08) marginally failed to reach significance.

**Figure 6.**
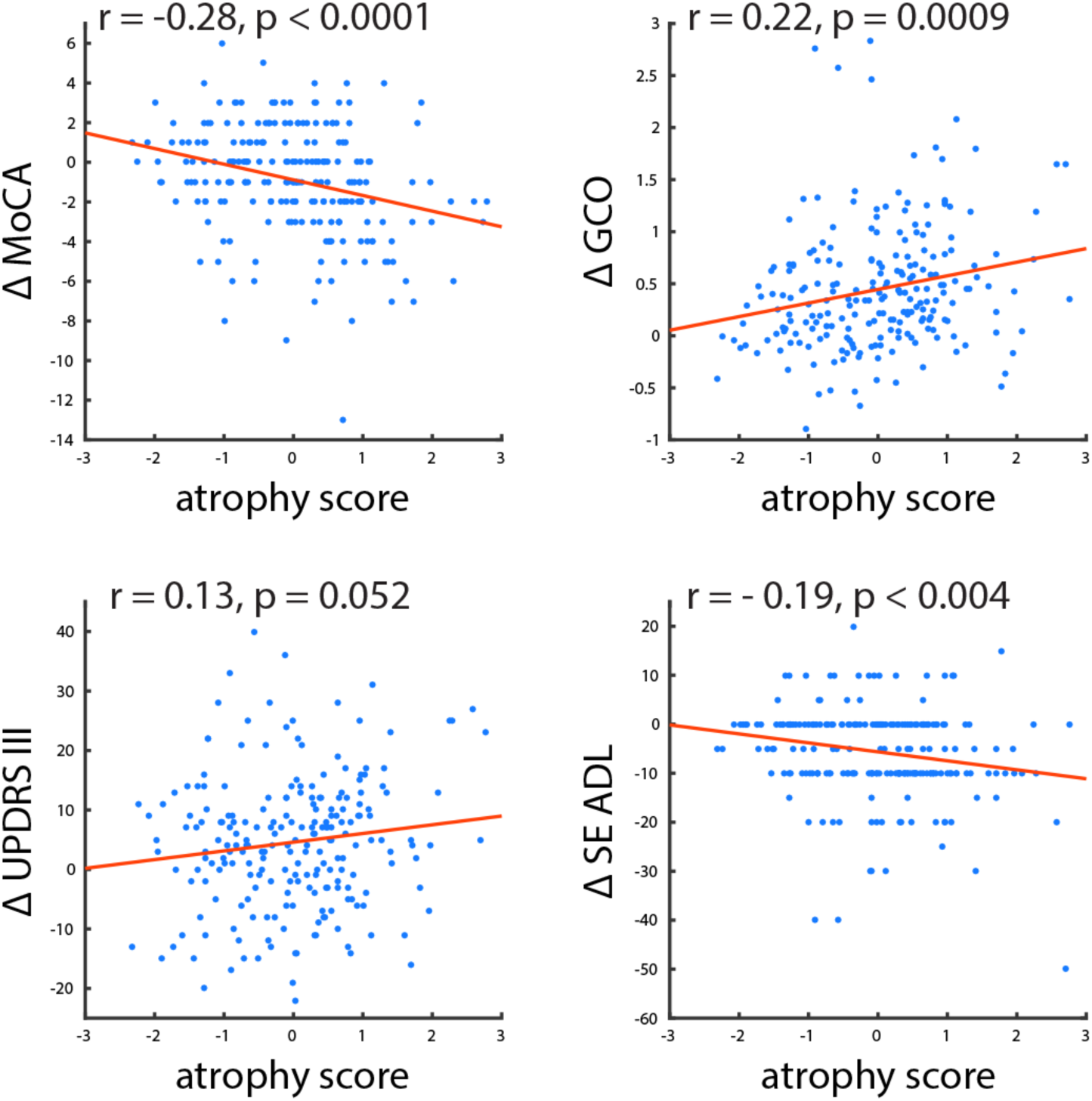
Baseline atrophy is associated with longitudinal clinical progression. Individual patients’ atrophy score (expression of the atrophy network from the PLS model) is correlated with longitudinal change in clinical measures of disease severity. PLS= Partial Least Squares. MoCA= Montreal Cognitive Assessment. GCO= Global Composite Outcome. UPDRS= Unified Parkinson’s Disease Rating Scale. SE ADL= Schwab and England ADL score (overall activities of daily living).

### 4. Discussion

The present study links multiple domains of clinical and biomarker features of PD to the underlying brain atrophy pattern using a single integrated analysis in a recently diagnosed population. In this *de novo* cohort, in addition to higher age, a wide range of motor and non-motor features were linked to brain atrophy. We hope the PLS approach used here provides a means to investigate the complex combination of motor and non-motor features of PD in relation to patterns of brain atrophy, as well as the intricate interplay between normal versus pathological aging.

Our findings suggest that a broadly distributed spatial pattern of brain atrophy is present in the early stages of PD, which covaries with motor, cognitive and other non-motor manifestations. This is somewhat at odds with the previous literature, where *de novo* PD is seldom associated with detectable brain atrophy. The participants in this cohort were all drug-naïve and within less than one year of diagnosis. A possible explanation for the greater ability of the multivariate approach to detect atrophy is that the course of PD may be stereotyped and the disease relatively widespread by the time early motor symptoms appear (Braak et al., 2003). Using all the voxels in the brain in a single analysis may confer greater sensitivity to deformation in a disease with a consistent spatial distribution. Although the first LV was associated with almost all the key clinical features of PD, we also describe two other LVs that capture smaller amounts of covariance (8-9% vs 17% for the first LV). These appear to respectively index a more benign clinical phenotype (tremor-dominant) with atrophy in motor areas and a more severe phenotype (postural instability – gait disorder) associated with brainstem and cortical atrophy. These patterns may indicate different potential modes of disease propagation, as evidenced in dementia using eigenvalue decomposition (Raj et al., 2012).

PD studies using brain imaging to date have almost always focused on differences between PD and healthy controls, or on a particular symptom manifestation (such as dementia) to study brain alterations. As a multivariate approach, PLS enables us to investigate brain alterations in PD subjects without a need for a control group and to consider multiple clinical aspects of the disease simultaneously.

We used our standard image analysis pipeline to calculate DBM as a measure of brain alterations. This pipeline (Aubert-Broche et al., 2013) has been previously used for several multicenter and multi-scanner studies and it has been shown to produce robust results by removing site-specific biases (Boucetta et al., 2016; Sanford et al., 2017; Zeighami et al., 2015). Also, in an earlier study, we provided evidence that DBM was a more sensitive measure of atrophy than VBM, especially for subcortical areas (Zeighami et al., 2015).

While the presence of atrophy early in the course of the disease is rarely reported, the direction of associations between atrophy and different clinical features and biomarkers is consistent with the literature. As one notable example, older age of onset and male gender were associated with greater expression of the PD-related pattern that was later demonstrated to correlate with faster progression. This is in line with previous reports of poorer prognosis of PD in older male patients (Post et al., 2011). Also, key non-motor features such as RBD, somnolence, autonomic disturbance and mood disorders contributed to the latent variable, consistent with the prognostic importance of these manifestations in other PD cohorts (Fereshtehnejad et al., 2015). Cognitive deficit, even though mild in severity, was also a significant correlate of brain atrophy. Although definite cognitive impairment was an exclusion criterion in PPMI, mild cognitive impairment still significantly correlated with the pattern of atrophy. Up to one fifth of the early PD populations meet the criteria for mild cognitive impairment, which is a strong predictor of earlier onset of dementia and poor prognosis (Pedersen et al., 2013, 2017). It is noteworthy that visuospatial and executive functioning more prominently contributed to the pattern of atrophy than the other cognitive domains. This is consistent with other studies of cognitive impairment in PD compared to Alzheimer’s disease (Watson and Leverenz, 2010; Wu et al., 2012). The patterns of brain atrophy and related motor, autonomic and cognitive deficits identified in LV-I are consistent with each other: autonomic and sleep dysfunction are explained by brainstem atrophy, and cognitive deficits in the domains of attention, memory, and executive function are consistent with the involvement of frontal lobes, medial temporal lobes, and posterior visual areas.

PD- and age-related brain alterations can happen independently, however, it is more plausible that aging and neurodegeneration interact (Collier et al., 2011). To distinguish between normative and pathological aging and their effects in our analyses, we regressed out the effect of normative aging – obtained from healthy subjects in the same dataset - from brain deformation maps of the PD patients. In contrast to the non-age-regressed results, two significant distinct patterns emerged. First, affective and sleep-related symptoms were more prominent contributors in the absence of any significant contribution of age and symptom duration. This aligns with the prodromal phase of PD during which the majority of the non-motor features reach high severity and precede the appearance of motor symptoms (Pfeiffer, 2016). The main exception is autonomic disturbance (SCOPA score and BP Sys drop), which usually worsens alongside PD progression. This is manifested in the second age-regressed LV, which may represent the pathological aging phenomenon in PD.

Even though, the effect of normative aging was regressed out in this complementary analysis, age and symptom duration remained as prominent contributors of the AR-LV-II atrophy pattern. Overall, this second age-regressed LV may represent pathological or accelerated aging in PD, as it also featured greater motor severity and more dopaminergic denervation (as measured by SPECT SBR).

Using PLS, we obtained a disease related atrophy map that included brainstem (medulla in the area of the dorsal motor nucleus of the vagus, red nucleus and substantia nigra), basal ganglia (including putamen, caudate, pallidum and subthalamic nucleus), cortical regions, as well as cerebellar regions. These findings are consistent with the earlier stages of Braak’s description of disease spread (Braak et al., 2003), as well as our previously published PD atrophy network map, based on this dataset (Zeighami et al., 2015). It is notable that atrophy was also identified in frontal regions, belonging to Braak Stage V (Braak et al., 2003), and not usually thought to be affected at the time of diagnosis. In that report, Braak et al. only noted frontal cortex Lewy pathology in patients at Hoehn and Yahr stage III or greater, which typically occurs at least 24-36 months after diagnosis (Zhao et al., 2010). This raises the possibility that brain atrophy may precede the arrival of synucleinopathy possibly due to tissue loss secondary to deafferentation.

One of the main strengths of the proposed approach is the ability to detect brain-clinical manifestations of the disease at an early stage. We further show that the PLS scores relate to disease progression in the follow up visits. These results provide an opportunity to develop a simple comprehensive measure per subject which can be used as a prognostic biomarker of the disease. This approach could also have value in assessing prodromal disease populations, identified through genetic testing or the presence of RBD. We suggest that it could also be applicable to other neurodegenerative or neurodevelopmental diseases.

The findings from this study should be considered in light of some limitations. Using PLS provides the opportunity to comprehensively investigate brain-clinical relations. However, we lose specificity as to how each particular clinical manifestation potentially relates to a specific brain region, rather than the atrophy pattern as a whole. Such individual relationships need to be addressed in future studies using independent PD cohorts. While we investigated the relationship between baseline findings and longitudinal clinical changes, future studies also need to investigate longitudinal brain alterations in PD and how they relate to disease progression.

In this study, we have taken advantage of PLS as a multivariate approach to investigate the collective relationship between brain alterations reflected in DBM measures and various aspects of the disease reflected in clinical measurements. We used data consisting of people with early diagnosed, drug-naïve PD who were followed for an average of 2.7 years from PPMI, a global multi-center study. While 2.7 years is a relatively short-term follow-up, the atrophy pattern was significantly associated with the longitudinal rate of decline in several clinical measures. In other words, high-scoring participants with more atrophic patterns at baseline experienced faster progression on the global single indicator of all symptom categories as well as the cognitive measure. Taken together, this study provides a new framework for studying neurodegenerative diseases with multi-faceted clinical measures and the interactions between brain alterations and disease manifestations. In addition, the single collective score summarizing the disease burden for each individual subject can be used as a potential biomarker for both diagnostic and prognostic purposes.

## Acknowledgement

This research was supported by the Canadian Institutes for Health Research, Natural Sciences and Engineering Research Council of Canada, Michael J Fox Foundation, Weston Brain Institute, and the Alzheimer’s Association. PPMI – a public-private partnership – is funded by the Michael J. Fox Foundation for Parkinson’s Research and funding partners, including AbbVie, Avid, Biogen, Bristol-Myers Squibb, Covance, GE Healthcare, Genentech, GlaxoSmithKline, Lilly, Lundbeck, Merck, Meso Scale Discovery, Pfizer, Piramal, Roche, Sanofi Genzyme, Servier, Teva, and UCB.

